# Pervasive Horizontal Transfer of Adeno-Associated Virus Capsid Genes

**DOI:** 10.1101/2025.03.15.643461

**Authors:** Robert J. Gifford

## Abstract

Adeno-associated viruses (AAVs) are non-pathogenic DNA viruses with potent gene delivery capabilities, making them essential tools in gene therapy and biomedical research. Despite their therapeutic importance, key aspects of AAV natural biology remain obscure, complicating efforts to explain rare AAV-associated diseases and optimize gene therapy vectors. By analyzing sequence data from virus isolates and endogenous viral elements (EVEs), I reveal a striking evolutionary pattern: while AAV sub-lineages, defined by the replication-associated (*rep*) gene, have broadly co-diverged with host groups over millions of years, capsid (*cap*) diversity has been shaped by extensive recombination. In particular, one capsid lineage, Mammalian-wide (*M-wide*), has spread horizontally across diverse rep lineages and host taxa through multiple recombination events. Furthermore, several AAVs with M-wide capsids - including AAV-4, AAV-12, and bovine AAV (BAAV) - originate from historical adenovirus (Ad) stocks, raising the possibility that laboratory conditions contributed to capsid transfer. Distinguishing natural from laboratory-driven recombination is essential for understanding AAV ecology and its implications for gene therapy. A systematic sequencing effort in human and primate populations is needed to assess the extent of recombinant capsid acquisition, determine the impact of laboratory-driven recombination on circulating AAV diversity, and track ongoing recombination events that could affect vector safety and efficacy.

## Introduction

Adeno-associated viruses (AAVs) are small, non-enveloped, single-stranded DNA viruses of the *Dependoparvovirus* genus (family *Parvoviridae*) [1]. They require co-infection with helper viruses, such as adenoviruses or herpesviruses, for productive replication [2]. The AAV genome encodes two primary genes: *rep* (essential for replication and genome packaging) and *cap* (encoding the structural proteins of the viral capsid).

AAVs are widely used in gene therapy, where the viral coding region is replaced with a therapeutic transgene, and the *rep* and *cap* genes are supplied in *trans*. Recombinant AAV (rAAV) vectors are now approved for treating genetic disorders such as hemophilia and inherited retinal diseases, and show growing potential across a broad range of therapeutic applications [3]. Capsid proteins, which determine host range and tissue specificity, are key targets for modification to improve gene delivery, transduction efficiency, and tropism, as well as to evade neutralizing antibodies from prior AAV exposure [4].

Despite decades of research on rAAV vectors, the natural biology of AAVs remains poorly understood. Like other parvoviruses, AAVs exhibit high recombination and mutation rates [5, 6], yet their extensive genomic ‘fossil record’, comprised of EVEs, reveals a remarkable degree of sequence conservation over millions of years [7]. Sequence and serological data indicate that diverse AAVs circulate in humans and non-human primates [5, 8], but virus-host interactions remain opaque. While AAVs are generally considered apathogenic [2], or even beneficial [9], recent reports linking AAV-2 to unexplained cases of childhood hepatitis [10], along with emerging evidence of context-dependent pathogenicity [11], highlight the gaps in our understanding of AAV-host interactions. Clarifying these relationships is critical not only for assessing the risks of naturally circulating AAVs but also for designing safer and more effective gene therapy vectors.

Here, I show that AAV capsids have undergone extensive horizontal transfer across divergent lineages, a process that may influence both viral evolution and gene therapy applications. These findings also raise critical questions about the role of laboratory-driven recombination in shaping AAV diversity.

## Results

Phylogenetic analysis of *rep* genes from AAV isolates and EVEs shows that AAVs form a distinct, well-supported subclade within genus *Dependoparvovirus*. Among AAVs that infect mammals, *rep*-based clades broadly align with host orders—Primates, Rodentia, Chiroptera, Artiodactyla— indicating long-term, stable host-virus associations. These ancient relationships are further supported by EVE data [7, 12] (**Fig. 1**).

**Figure 1.**
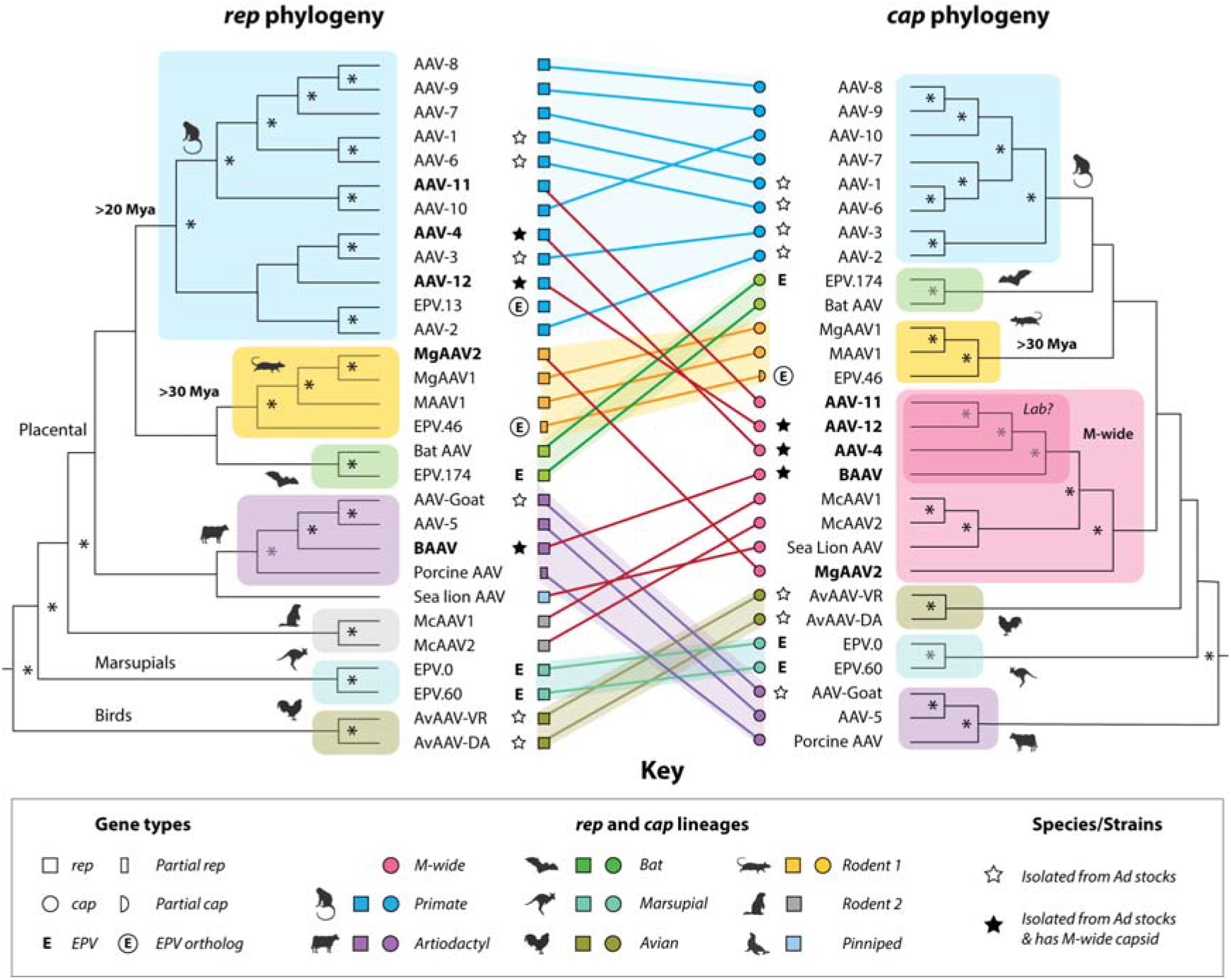
Tanglegram showing phylogenetic discordance between *rep* and *cap* and spread of the M-wide capsid lineage. Comparison of *rep* (left, outgroup-rooted) and *cap* (right, midpoint-rooted) phylogenies of adeno-associated viruses (AAVs), illustrating recombination. Shaded clades correspond to host taxonomic groups, as indicated in the key. Connecting lines highlight rep/cap discordance and suggest independent acquisitions of the M-wide capsid (pink), found across multiple mammalian orders. Bold labels mark taxa potentially acquiring M-wide. A darker pink region indicates a sub-lineage of M-wide AAVs associated with adenovirus (Ad) stocks. Endogenous parvoviral elements (EPVs) are marked by symbols (see key); orthologous EPVs provide minimum age estimates (rounded to nearest 5 My) at the base of the respective clades—e.g., *EPV-dependo*.*13-cercopithecidae* (>20 My) and *EPV-dependo*.*46-gliridae* (>30 My). Asterisks indicate bootstrap support >70%.

In contrast, *cap*-based phylogenies reveal a markedly different topology. While most *cap* lineages align with mammalian host orders, one prominent exception stands out: a single capsid lineage – here termed Mammalian-wide-1 (*M-wide*) – appears across multiple *rep* lineages, consistent with widespread horizontal transfer via recombination (**Fig. 1**). These incongruent relationships are supported by statistical comparison of tree topologies (Shimodaira–Hasegawa test) and orthogonal recombination analyses, including GARD and split network methods (see **Methods**).

Strikingly, three AAVs bearing M-wide capsids – AAV-4, AAV-12, and bovine AAV (BAAV) – were originally isolated from adenovirus stocks established in the 1950s (Fig. 1, Table 1). Their *cap* genes form a well-supported subclade (**Fig. 1**), suggesting a shared origin potentially linked to laboratory culture.

**Table 1.** Isolation History & Host Associations of Primate and Artiodactyl Adeno-Associated Viruses.

| AAV Isolation history |  |  | Ad Isolation history |  |  |  |  | AAV-Host Associations |  |  |
| --- | --- | --- | --- | --- | --- | --- | --- | --- | --- | --- |
| Name<sup>1</sup> | Iso. Source<sup>2</sup> | Year | Ad Strain(s) | Iso. Source | Cell culture | Year<sup>3</sup> | Ad Host | Primary Host<sup>4</sup> | Seroprevalence<sup>5</sup> |  |
|  |  |  |  |  |  |  |  |  | Human | NHP |
| AAV-1 | Ad stocks† | 1965 | Ad7 | Human | MK, Human | 1953 | *H. sapiens* | Humans | ++++ | +++ |
| AAV-2 | Ad stocks† | 1965 | Ad12, Ad7 | Human | Human | 1953 | *H. sapiens* | Humans | +++++ | ++ |
| AAV-3 | Ad stocks† | 1965 | Ad7 | Human | AGMK | 1953 | *H. sapiens* | Humans | +++ | ++ |
| AAV-4* | Ad stocks | 1968 | SV15 | Macaque | MK, AGMK | 1956 | *M. mulatta* | NHP | + | +++++ |
| AAV-5 | Hu. Tissue | 1982 | *n/a* |  |  |  |  | Humans | +++ | ++ |
| AAV-6 | Ad stocks | 1998 | Ad5 | Human | Human | 1953 | *H. sapiens* | Humans | ++++ | ++ |
| AAV-7 | OWM Tissue† | 2004 |  |  |  |  |  | NHP | ++ | +++++ |
| AAV-8 | OWM Tissue† | 2004 |  |  |  |  |  | NHP | ++ | +++++ |
| AAV-9 | Hu. Tissue† | 2004 |  |  |  |  |  | NHP, Humans | +++ | +++++ |
| AAV-10 | OWM Tissue | 2004 |  |  |  |  |  | NHP | +++ | +++++ |
| AAV-11* | OWM Tissue | 2004 |  |  |  |  |  | NHP | n/k | ++ |
| AAV-12* | Ad stocks | 2008 | SV18 | Vervet | n/k | *n/k* | *C. aethiops* | NHP | - | n/k |
| BAAV* | Ad stocks† | 1970 | Bovine Ad | Cow | Bovine | 1957 | *B. taurus* | Cattle | n/k | n/k |
| AAV-Go | Ad stocks | 2004 | Caprine Ad | Goat | Caprine | 2004 | *C. hircus* | Goats | n/k | n/k |

## Discussion

AAVs were first identified in the 1960s as contaminants of adenovirus (Ad) stocks [13]. These stocks – which were derived from primate and bovine tissues—had been established a decade earlier, in an era when few viruses could be propagated outside live hosts. Given the limited virological tools available, AAVs likely remained undetected in early adenovirus cultures for years. These early Ad stocks were often derived from pooled or heterogeneous biological samples, creating an environment conducive to AAV co-infection. The presence of helper viruses, elevated viral titers, and permissive cell lines may have enabled capsid gene exchange through recombination, and inadvertently contributed to the emergence and spread of recombinant AAVs within laboratory cultures.

Since the 2000s, AAVs encoding M-wide capsids have been detected in both primate centers and dairy cattle populations [15]. If these virus lineages did originate in laboratory cultures, they now appear to be established in natural hosts. Notably, one such virus (AAV-11) is >99% identical in the *rep* gene to AAV-10, which encodes a canonical primate AAV capsid, suggesting that recent recombination events continue to shape AAV diversity. If capsid exchange is ongoing, the functional breadth of M-wide capsids in primate AAVs, spanning epithelial, intestinal, and lymphoid tropisms, implies that such recombination may have significant biological and therapeutic implications [3, 4].

While laboratory propagation may have played a role in M-wide’s spread, an alternative possibility is that this capsid lineage was already naturally widespread, with its presence in AAVs derived from Ad stocks reflecting only pre-existing bovine and primate AAV diversity, rather than lab-driven recombination. Disentangling these scenarios is key to understanding AAV ecology and evolution.

Modern sequencing technologies can facilitate broad-scale characterization and surveillance of AAV diversity. Such efforts would not only clarify the natural ecology and evolution of AAVs but also inform gene therapy applications by identifying circulating diversity and potential immune interactions.

## Materials and Methods

AAV genome and EVE sequences were obtained from public databases and curated in a reproducible database framework integrating standardized alignments and metadata [12, 16]. Phylogenetic trees were reconstructed using maximum likelihood from conserved *rep* and *cap* coding regions. Recombination was assessed by comparing tree topologies and applying complementary analytical approaches. All curated sequences, alignments, and analysis workflows are fully documented and openly accessible in the associated repositories, and a Docker image is provided for streamlined, cross-platform reproducibility [12].

## Acknowledgment

I thank Professor Robert Kotin for his valuable feedback.

