## Extended Methods for "Pervasive Horizontal Transfer of Adeno-Associated Virus Capsid Genes"

**Sequence Data and Classification**

AAV genomic sequences were retrieved from GenBank and curated in AAV-Atlas, a GLUE-based platform integrating standardized metadata, alignments, and serotype classifications [1]. To focus on naturally circulating AAV diversity, sequences associated with patents or laboratory-derived constructs were excluded. Classification was performed using Maximum Likelihood Clade Assignment (MLCA) in GLUE [2]. AAV-derived EVEs were sourced from Parvovirus-GLUE-EVE, a repository cataloging viral loci with genomic context and orthology relationships [3]. EVE orthology across host species was confirmed using comparative genomic analysis of flanking regions via BLASTn and whole-genome alignment.

**Gene Identification and Alignment**

The *rep* and *cap* coding regions were extracted based on AAV-Atlas reference genome annotations [1]. Codon-aware alignment was performed in GLUE. Multiple-sequence alignments were generated using MUSCLE [4], and manually curated in Se-Al [5], ensuring reading frame integrity for EVEs via virtual translation.

**Definition of Phylogenetic Partitions**

To improve phylogenetic signal and minimize alignment noise, conserved coding regions in the *rep* and *cap* genes were subdivided into well-defined partitions. These were selected manually, based on conservation patterns and indel rates observed using Se-Al [5] and computed using GLUE’s ‘amino acid frequency’ function [2]. Partition boundaries were chosen to align with:

- Highly conserved amino acid motifs (typically ≥90% identity; often invariant),
- Regions with low observed indel frequency across taxa, and
- Maintenance of codon structure in the reference-constrained alignment (AAV2; AF043303).

**Phylogenetic Reconstruction**

Maximum likelihood phylogenies were inferred separately for *rep* and *cap* genes using RAxML (v8.2.12) [6], with optimized partitioning to maximize taxon representation. The *rep* tree was rooted using *Dependoparvovirus* outgroups, while the *cap* tree was midpoint-rooted. Bootstrap support (1,000 replicates) was calculated under the GTR+G model for nucleotide sequences and the JTT model for amino acid sequences, selected via likelihood ratio testing.

**Phylogenetic Incongruence Testing**

Phylogenetic incongruence between rep and cap genes was assessed using the Shimodaira-Hasegawa (SH) test in CONSEL [7]. Maximum likelihood trees were inferred separately for *rep* and *cap* using RAxML (v8.2.12) [6], and per-site log-likelihoods were calculated on a concatenated alignment containing high-confidence sites. The Approximately Unbiased (AU) test strongly rejected the rep tree (AU = 0.006, SH = 0.012) in favor of the *cap* tree (AU = 0.994, SH = 0.988), supporting independent evolutionary histories shaped by recombination.

**SplitsTree Analysis**

To assess phylogenetic compatibility independent of explicit tree reconstruction, we generated a splits network using SplitsTree5. The nucleotide alignment included all conserved partitions (Rep78 parts 1–5 and VP1 parts 1–7), exported from GLUE using the alignmentColumnsSelector module. The resulting nexus file was analyzed using default NeighborNet settings under uncorrected p-distances, with taxa labeled by serotype or clade. Reticulation in the output network was interpreted as evidence of recombination.

**GARD Analysis**

We submitted the same partitioned alignment described above for SplitsTree analysis to GARD (Genetic Algorithm for Recombination Detection), using the Datamonkey web server. GARD performs model-based inference of recombination breakpoints by evaluating phylogenetic incongruence between partitions. The analysis identified twelve statistically supported breakpoints, which closely tracked the boundary regions between the conserved partitions defined in our tree-based analysis. Full output is provided in the AAV-Atlas repository archived in Zenodo [1].

**Estimation of Divergence Time**

Pairwise sequence divergence was calculated within the rep gene between AAV-10 and AAV-11, spanning 1,869 nucleotides. The two viruses differed at 9 positions (0.482% sequence divergence). A molecular clock model using published substitution rates for single-stranded DNA viruses [8], ranging from 1×10⁻⁵ to 2×10⁻⁴ substitutions/site/year, estimated a divergence time between 12 and 241 years ago.

**Serological Data Compilation**

To estimate the relative prevalence of neutralizing antibodies (NAbs) against AAV serotypes in humans and non-human primates, I conducted a structured review of published serological studies. My goal was to provide a qualitative summary of general trends across serotypes that could be used for comparative purposes in **Table 1**. The analysis was not intended to be exhaustive or to yield definitive epidemiological estimates, but rather to support interpretation of AAV-host associations in phylogenetic context.

I reviewed 11 primary studies and reviews that reported IgG or NAb data for multiple AAV serotypes across human or non-human primate cohorts. These included historical serosurveys conducted before the development of modern molecular tools, such as those by Blacklow et al. (1968, 1971) [9, 10] and Sprecher-Goldberger et al. (1971) [11], as well as more recent studies using standardized neutralization assays and broader sampling, such as those by Boutin et al. (2010) [12], Calcedo et al. (2009) [13], and Klamroth et al. (2022) [14]. Relevant findings were also drawn from synthesis papers and methodological comparisons.

Seroprevalence estimates were curated and harmonized across sources using a six-point categorical scale (– / + / ++ / +++ / ++++ / +++++) to reflect approximate antibody levels while accounting for differences in assay sensitivity, neutralization thresholds, and cohort size. Where studies disagreed, I prioritized those with broad geographic coverage, internally consistent methodology across serotypes, and clear reporting of cohort composition. In general, data from Calcedo et al. [13] and Boutin et al. [12] were given greater weight due to their comprehensiveness and use of standardized neutralization assays.

**Data Availability**

All sequence alignments, phylogenetic trees, and analysis pipelines are publicly available in the AAV-Atlas repository [1]. This repository provides platform-independent tools and datasets for full reproducibility of the analyses described in this study.
